# Viral diversity, ecological interconnectedness, and the identification of mammalian chuviruses in Australian microbats

**DOI:** 10.1101/2025.11.29.691345

**Authors:** Ayda Susana Ortiz-Baez, Julien Mélade, Kate Van Brussel, Bethan J. Lang, Anna Lachenauer, Edward C. Holmes

**Affiliations:** School of Medical Sciences, University of Sydney, Sydney, New South Wales 2006, Australia; Department of Dermatology, Stanford University School of Medicine, Redwood City, CA, USA

## Abstract

Microbats are a large and ecologically important group of Australian mammalian fauna. However, their RNA virome diversity, as well as its ecological and evolutionary significance, has received limited study. We applied a metatranscriptomic approach to reveal more of the diversity of RNA viruses present in faeces from different microbat species in New South Wales and South Australia, including the critically endangered Southern bent-wing bat (*Miniopterus schreibersii bassanii*) from the Naracoorte bat maternity caves in South Australia. The data generated revealed a high diversity of RNA viruses, including 51 likely mammalian-associated viruses classified into ten taxonomic groups, including the *Coronaviridae*, *Hepeviridae* and *Chuviridae*. Notably, we identified a mammalian-specific lineage of chuviruses associated with bats in Australia and of bats and rodents in China, strongly suggesting that viruses of this family have established sustained transmission cycles in mammals as well as invertebrates. Our results also revealed widespread viral connectivity among alphacoronaviruses across multiple microbat species in mainland Australia and Christmas Island, indicative of long distance viral movement. High viral diversity and virus co-circulation was observed within the Southern bent-wing bat population of the Naracoorte caves, suggesting complex population dynamics that might facilitate virus maintenance and transmission. Overall, these findings highlight the role of Australian microbats as viral reservoirs, including the presence of viruses not previously associated with sustained mammalian transmission.

## Introduction

Comprising over 1,400 species, Chiroptera (i.e., bats) is one of the most species-rich orders of mammals, ranking second only to Rodentia [1]. This diversity encompasses a wide range of biological features, including varied diets, body sizes, sensory adaptations, hibernation patterns and roosting behaviours. Microbats, formerly classified as Microchiroptera, are now incorporated into the suborders Yinpterochiroptera and Yangochiroptera, and comprise small-sized, laryngeal echolocating bats [2]. These animals mainly rely on echolocation for navigating and foraging for insects, although diverse feeding behaviours are found in some species [1]. Prey composition primarily comprises lepidopterans, dipterans, as well as hemipterans and coleopterans among other arthropod taxa [3, 4].

Microbats roost in a wide range of habitats, including caves, crevices, and tree cavities [5]. To mitigate habitat loss and human-wildlife interactions, artificial roosting sites such as bat boxes are built to provide shelter to bats in urban and natural settings [5–7]. The occupancy of bat boxes by different species may vary depending on geographic location, seasonal dynamics, and the structural design of the boxes [7]. Compared to natural habitats, bat boxes may provide less effective insulation and hence result in increased ectoparasite accumulation [8, 9]. Depending on environmental conditions and food availability, microbats are also able to reduce their physiological activity and energy demand by entering torpor, especially as a mechanism to cope with low temperatures [5, 10]. Natural habitats such as caves provide shelter with more stable levels of temperature and humidity, which are essential for entering topor and breeding. For instance, maternity caves are well-suited for resting and nursing newborn bats, thereby supporting colony social structure, and hence contribute to bat conservation and population well-being.

Australia is home to 77 species of bats. While these species vary in their spatial distribution, some exhibit a wide range across the country. For example, the lesser long-eared bat (*Nyctophilus geoffroyi*) and Gould’s wattled bat (*Chalinolobus gouldii*) are endemic to Australia and occur in a wide range of habitats and environments [11]. In contrast, species like the large bent-wing bat (*Miniopterus schreibersii oceanensis*) and the little forest bat (*Vespadelus vulturnus*) are restricted to south-eastern Australia. These microbats exhibit also diverse roosting strategies, ranging from solitary behaviour to the formation of large seasonal colonies, with up to 35,000 individuals in the case of the Southern bent-wing bat (*Miniopterus orianae bassanii*) [12]. Bat congregation is often associated with reproduction, hibernation, mating, or the establishment of maternity colonies.

Many microbat species are experiencing a decrease in population size due to habitat loss, climate change, limited resource availability, and the presence of invasive species [13–15]. Population declines have been documented in Australia, including in the little forest bat, the white-striped free-tailed bat (*Tadarida australis*) and the yellow-bellied shear bat (*Saccolaimus flaviventris*), with the cave-roosting Southern bent-wing bat is considered a threatened species [16, 17]. These declines are largely due to habitat destruction caused by urban development, logging, mining, wildfires and the introduction of invasive species [11, 18].

Revealing the diversity of viruses in wild microbats provides important information on virus population structure and virus-host associations, and assists in the identification of viruses that may have implications for bat conservation as well as those with zoonotic potential [19, 20]. The most common RNA virus families reported in microbats are the *Coronaviridae*, *Rhabdoviridae, Astroviridae, Picornaviridae* and *Paramyxoviridae* [19, 21]. As bat conservation is paramount, documenting the bat virome represents an important way to identify potential pathogens that could impact the well-being of bat populations. Fortunately, faecal sampling offers a viable, non-invasive approach for the surveillance of virus with relevant tropisms, enabling the detection of both mammalian-origin viruses and diet-derived viruses shed in bat droppings [22, 23].

Although Australia harbours a remarkable diversity of microbats, the composition, evolution, and ecology of their mammalian-associated RNA virome is largely unexplored. Herein, we use a meta-transcriptomics (i.e., total RNA sequencing) approach to analyse faecal samples collected from microbats in natural and artificial roosting sites in New South Wales and South Australia. From the data generated we assess the RNA virus diversity, ecological patterns and virome connectivity among microbat populations.

## Methods

### Ethical approval and scientific licenses

The collection of bat faecal samples was conducted under approval from the University of Sydney Animal Ethics Committee (Project number: 2023/AE002279). Fieldwork in South Australia was performed under the relevant permit issued by the Government of South Australia, Department for Environment and Water (Permit number: E27285-3). Methods were designed to be not invasive and minimize disturbance to the animals and habitats.

### Sample collection

Faecal samples from microbat species were opportunistically collected by placing tarps under roosting sites, including bat boxes, in various locations in New South Wales (Fairfield), and South Australia (Adelaide), Australia. Additional faecal samples were collected from the entrance of the Naracoorte bat maternity caves in south-eastern South Australia (**Table 1**, **Figure 1**). To minimize disturbance to the animals and ensure their welfare, all samples were collected during spring, after the hibernation period. All samples were preserved in DNA/RNA Shield and stored at -80°C to aid RNA integrity and deactivate viruses. When microbat species identity was unknown, taxonomic classification was performed based on the presence of the mitochondrial COI marker gene in the meta-transcriptomic data, queried in the BOLD Identification System [24], and compared with the species reported for the visited locations, where available. The species identified in this study included the Eastern bent-wing bat (*Miniopterus schreibersii oceanensis*), little forest bat (*Vespadelus vulturnus*), the lesser long-eared bat (*Nyctophilus geoffroyi*), and the Southern bent-wing bat (*Miniopterus schreibersii bassanii*) (**Table 1**).

**Figure 1.**
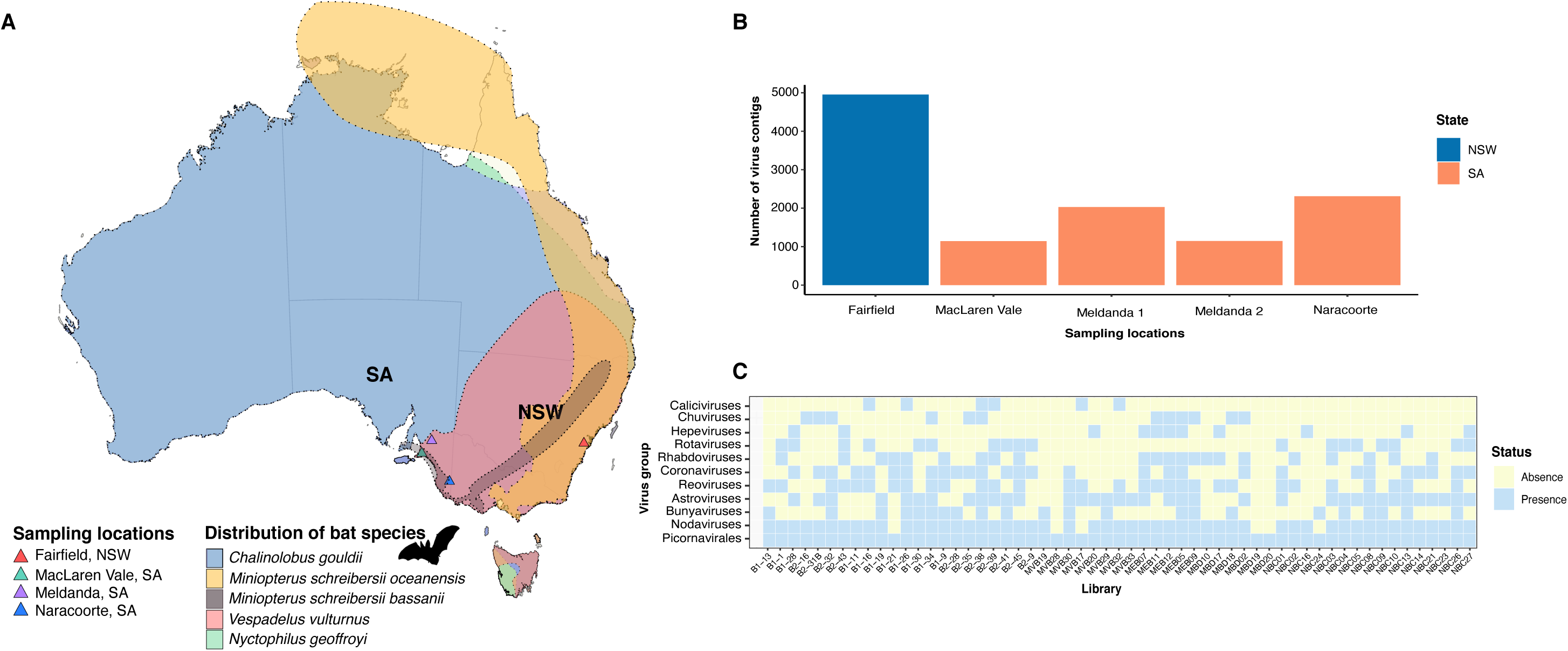
Overview of bat host range, viral contig counts per location and virus prevalence in the bat samples taken. (A) Range distribution of the most common bat species found in NSW and SA, Australia. Each species distribution is displayed in a different colour. Sampling locations are indicated with colour coded triangles. Species distributions were accessed from the IUCN and ATLAS web portals [11, 71]. (B) Number of virus contigs by sampling location. (C) Distribution of virus groups across the sample set. Each square denotes a single sample. Blue indicates virus presence, yellow denotes absence.

**Table 1.**
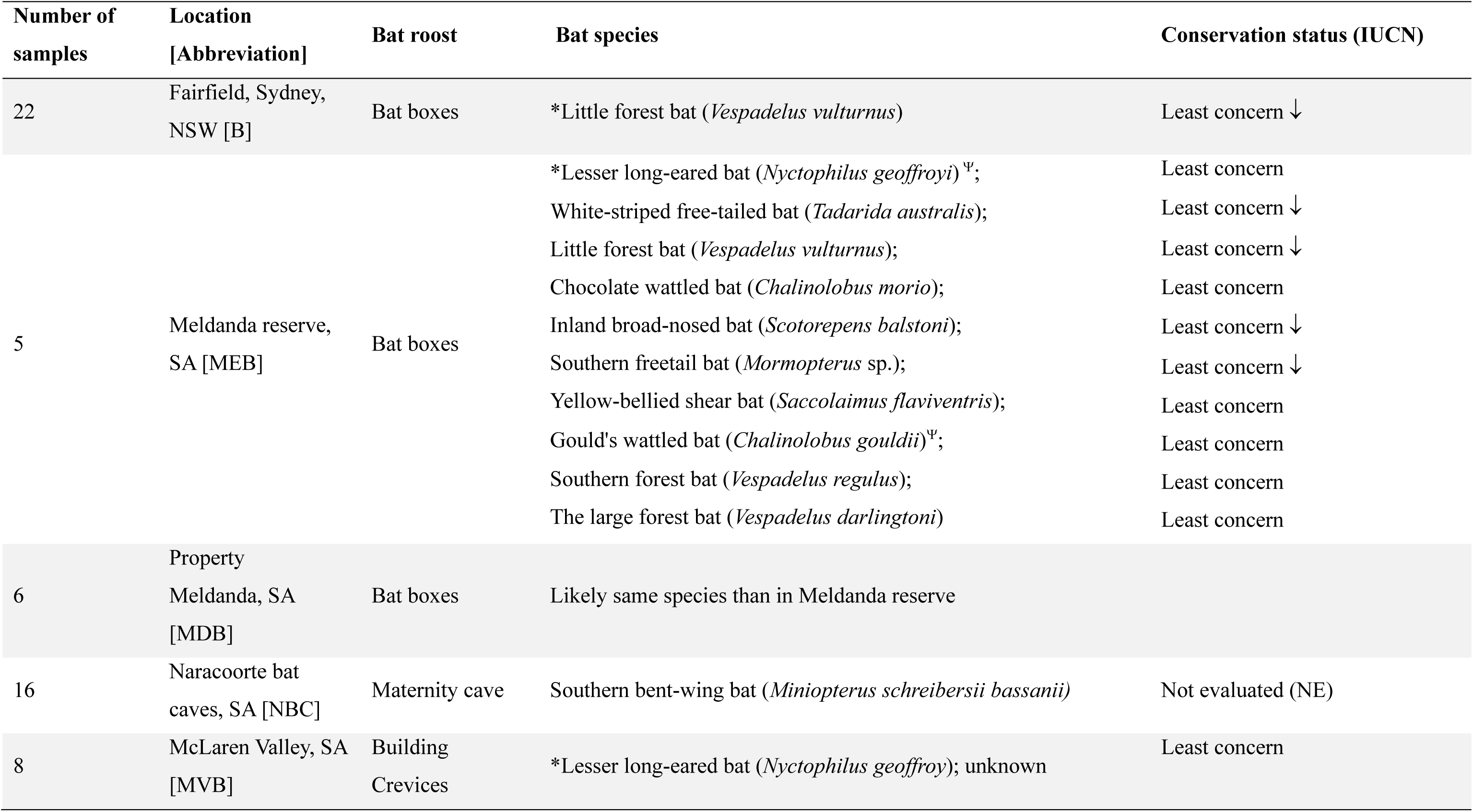
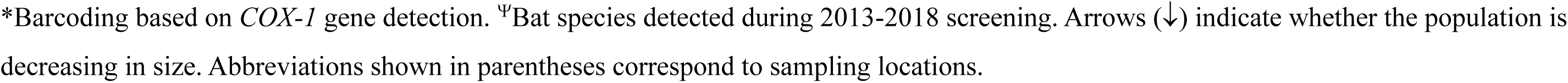
Bat sampling locations in New South Wales (NSW) and South Australia (SA), with the expected bat species at each site.

### Sample processing and sequencing

Bat faecal samples were thawed and homogenized by vortexing in the lysis reagent DNA/RNA shield at high speed for 30 seconds to ensure thorough mixing and shearing of the material. To facilitate the release of the nucleic acids and remove cellular debris, we used QIAshredder columns by centrifuging the samples at 4°C at maximum speed for 3 minutes. Total RNA was extracted using the QIAamp Viral RNA Kit (Qiagen, AU) according to the manufacturer’s instructions and quantified using the Qubit fluorometer.

The extracted RNA was subjected to ribosomal depletion, reverse-transcribed into cDNA, and then used as input for paired-end library preparation using the Illumina Stranded Total RNA Prep with Ribo-Zero Plus (Illumina). Finally, paired-end libraries were sequenced on the Illumina NovaSeq X platform using the 300 cycle.

### Raw data processing and assembly

Paired-end raw data were preliminary assessed with FASTQC v.0.11.8 [25] to identify low-quality ends and the presence of adapter sequences. Subsequently, reads were trimmed to maintain a Phred score of 25. A second round of quality assessment was then performed with FASTQC. Contigs were *de novo* assembled using Megahit v.1.29 with default settings [26]. Relative contig abundance was calculated as the number of transcripts per million (TPM) and reads per million (RPM) using RSEM v1.3.0 [27] and Salmon v.0.8.2 [28]. Cross-contamination between libraries due to index-hopping was considered likely when abundance values fell below 0.1% of the highest count recorded for a given contig across libraries.

### Contig annotation and taxonomic profiling

To identify potential virus sequences, contigs were initially screened against a database of RNA dependent RNA polymerase (RdRp) sequences using the RdRp-Scan tool [29]. These sequences were then compared against the nucleotide (NCBI-nt) and non-redundant (NCBI-nr) databases for taxonomic annotation and curation with e-value thresholds set to ≤ 1E-10 and ≤ 1E-4, respectively. Other virus gene fragments were identified using the protein Reference Viral DataBase (RVDB-prot) v28 database [30]. Open reading frames (ORFs) were predicted using the EMBOSS GetOrf tool [31], while domains and motifs were identified by scanning against all the Interpro databases using InterProScan v.5.63-95.0 [32]. The taxonomic composition of libraries was assessed using Kraken v.2.1.1 (core-nt database v.04-2024) [33] and MetaPhlan v.4.1.0 [34]. To identify clusters of sequences originating from different geographic regions, amino acid contig sequences were clustered using CD-Hit v.4.8.1, with a sequence identity threshold of ≥ 98% to group highly similar sequences.

### Abundance and phylogenetic analysis

The phylogenetic relationships of the virus sequences identified in the microbat data in the context of related sequences publicly available on the NCBI/GenBank database were estimated using the maximum likelihood (ML) method available in IQ-TREE v.2.3.6 [35]. In each case, the best-fit models of amino acid substitution for the RdRp sequences were assessed with the option -m TEST (**Supplementary Table S1**). Node support was estimated using the Shimodaira-Hasegawa approximate likelihood ratio test (SH-aLRT) and ultrafast bootstrap (UFBoot) support, with topological confidence set as SH-aLRT >= 80% and UFboot >= 95%, respectively. The likely host assignment for each virus, particularly whether they likely infected vertebrate or invertebrates (the latter being diet associated), was inferred from their phylogenetic relationships (i.e., vertebrate-associated viruses tend to form distinct clades), supplemented by metadata on sample origin (e.g., host, tissue type, geographic location), when available.

## Workflow and code availability

Most of the steps in this analytical procedure were followed as implemented in the Batch Artemis SRA Miner pipeline v. v1.0.4 (https://github.com/JonathonMifsud/BatchArtemisSRAMiner/tree/v.1.0.4), with adaptations made as needed.

## Results

### Mammalian-associated virus diversity in Australian microbats

To better understand the diversity and evolution of RNA viruses circulating in microbats in Australia, we sequenced 57 microbat faecal samples collected from several locations in New South Wales and South Australia (**Table 1**). When such information was available or based on direct observation of individuals, samples were taken from populations reported as apparently healthy. Sequencing yielded between 36 million and 312 million paired-end reads per library. A total of 3,192,497 contigs were assembled, including 11,580 of viral origin (**Figure 1**). Virus abundance varied between 0.01-215530 TPM.

Despite the small sample size we identified a broad range of likely mammalian- and diet-associated viruses spanning nine taxonomic groups: the orders *Picornavirales* and *Bunyavirales*, and the families *Nodaviridae, Hepeviridae*, *Coronaviridae*, *Astroviridae*, *Chuviridae*, *Rhabdoviridae*, *Reoviridae* and *Astroviridae* (**Figure 1**). Of these, picornaviruses were the most prevalent group in bat faecal samples (57/57 libraries), whereas hepeviruses and chuviruses were the least commonly detected (12/57 libraries) (**Figure 1**). Of note, 30 newly discovered viruses were identified in cave-roosting bats, while 95 newly discovered viruses were present in faecal samples collected from bat boxes. Approximately, 48.8% of the viruses discovered (n = 61) were likely mammalian in origin in that they were related to other mammalian-associated viruses on phylogenetic trees. Prior to formal classification, we provisionally named the newly discovered viruses as “AusMicrobat” viruses.

The bat viruses identified here that were most closely related to known viruses belonged to the *Coronaviridae* (99.5% amino acid identity to known viruses in the RdRp) and the *Sedoreoviridae* (76–99% amino acid identity to known viruses in the VP1) (**Table 2**). The novel astroviruses exhibited moderate to high similarity to other bat astroviruses (∼60–93% amino acid identity in the capsid), while some had similarity to avian astroviruses (∼53% amino acid identity). The most genetically divergent virus sequences were also identified in the *Sedoreoviridae*, including AusMicrobat sedoreoviruses 1 and 8 (23 – 36% identity to known viruses), and were likely diet-associated as they were unrelated to known vertebrate viruses. Viral abundance estimates indicated that reoviruses (TPM = 0.76–24225) and picornaviruses (TPM = 0.16–2200) were relatively abundant, while rhabdoviruses were the least abundant (TPM = 0.52–5.84). Also of note was that the viruses identified in the bat faecal samples collected from SA and NSW were often closely related. For example, highly similar coronaviruses (up to 96% genome-scale amino acid identity) and astroviruses (up to 93% amino acid identity in the capsid protein) were detected between both states, and which also clustered phylogenetically with viruses found in various locations in Australia as well as Christmas Island (a remote Australian territory located about 2600 km from the mainland) (**Figure 2**, **Figure 3**).

**Figure 2.**
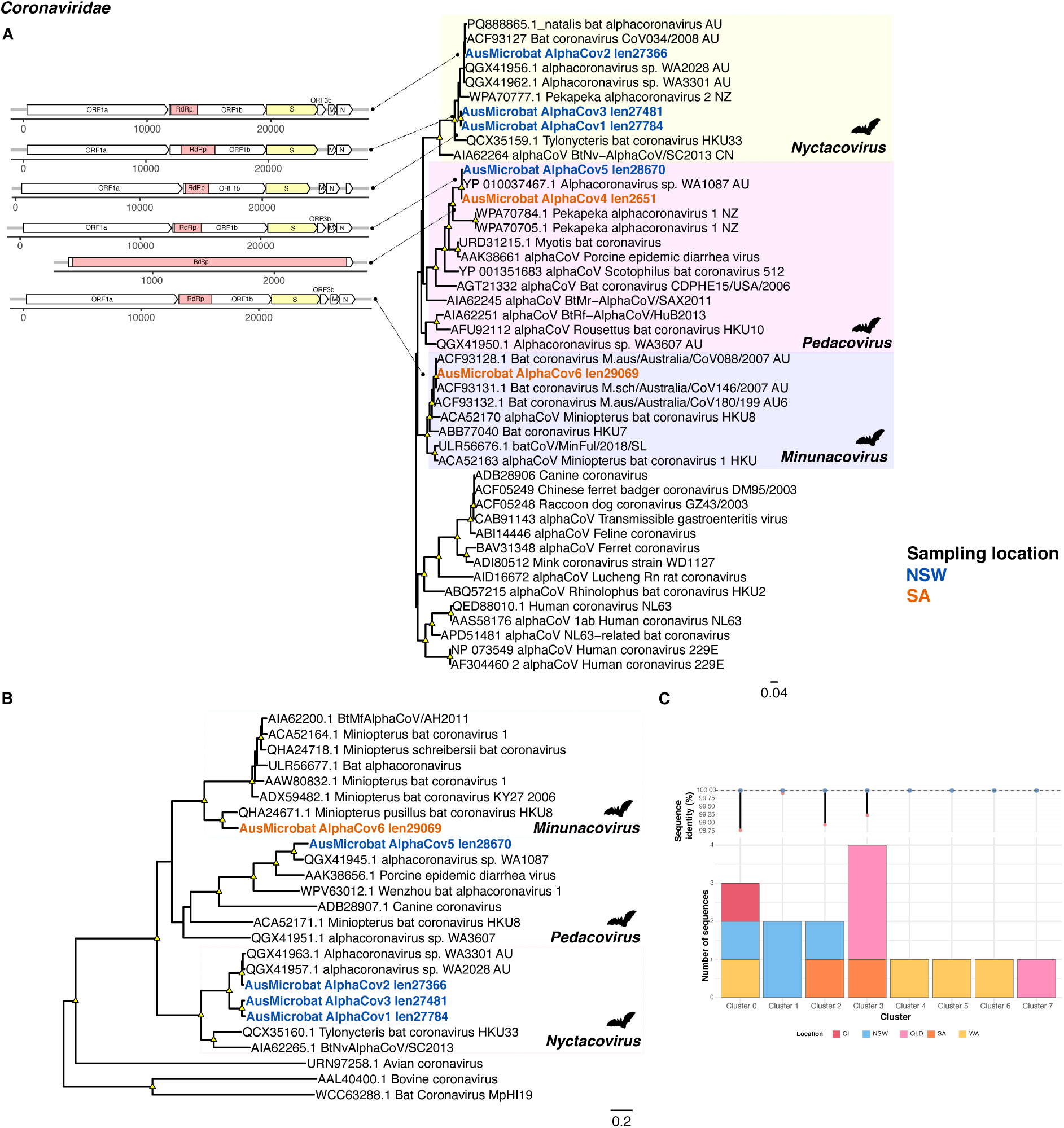
Phylogenetic relationships among the alphacoronaviruses identified in this study. Maximum likelihood trees are based on the amino acid sequences of the (A) RdRp – nsp12 and (B) Spike protein. All phylogenetic trees are mid-point rooted for clarity only. Scale bars represent the number of amino acid substitutions per site. Known genera are represented by colour-coded clades for easier interpretation. Tip labels for novel viruses are coloured by sampling location – blue (NSW), orange (SA). Nodal support values ≥80% SH-aLRT and ≥95 % UFboot are denoted with yellow triangles at nodes. Bat-associated lineages are indicated with bat silhouettes. A schematic representation of the newly discovered alphacoronavirus sequences is displayed next to the tip labels, with annotated ORFs and the RdRp and spike domains shown in colour. (C) Clustering of alphacoronavirus sequences across multiple sampling locations in Australia. Each cluster comprises sequences sharing ≥ 98% amino acid sequence identity with the representative sequence. Australian locations are indicated as follows: CI (Christmas Island), NSW (New South Wales), WA (Western Australia), QLD (Queensland) and SA (South Australia).

**Figure 3.**
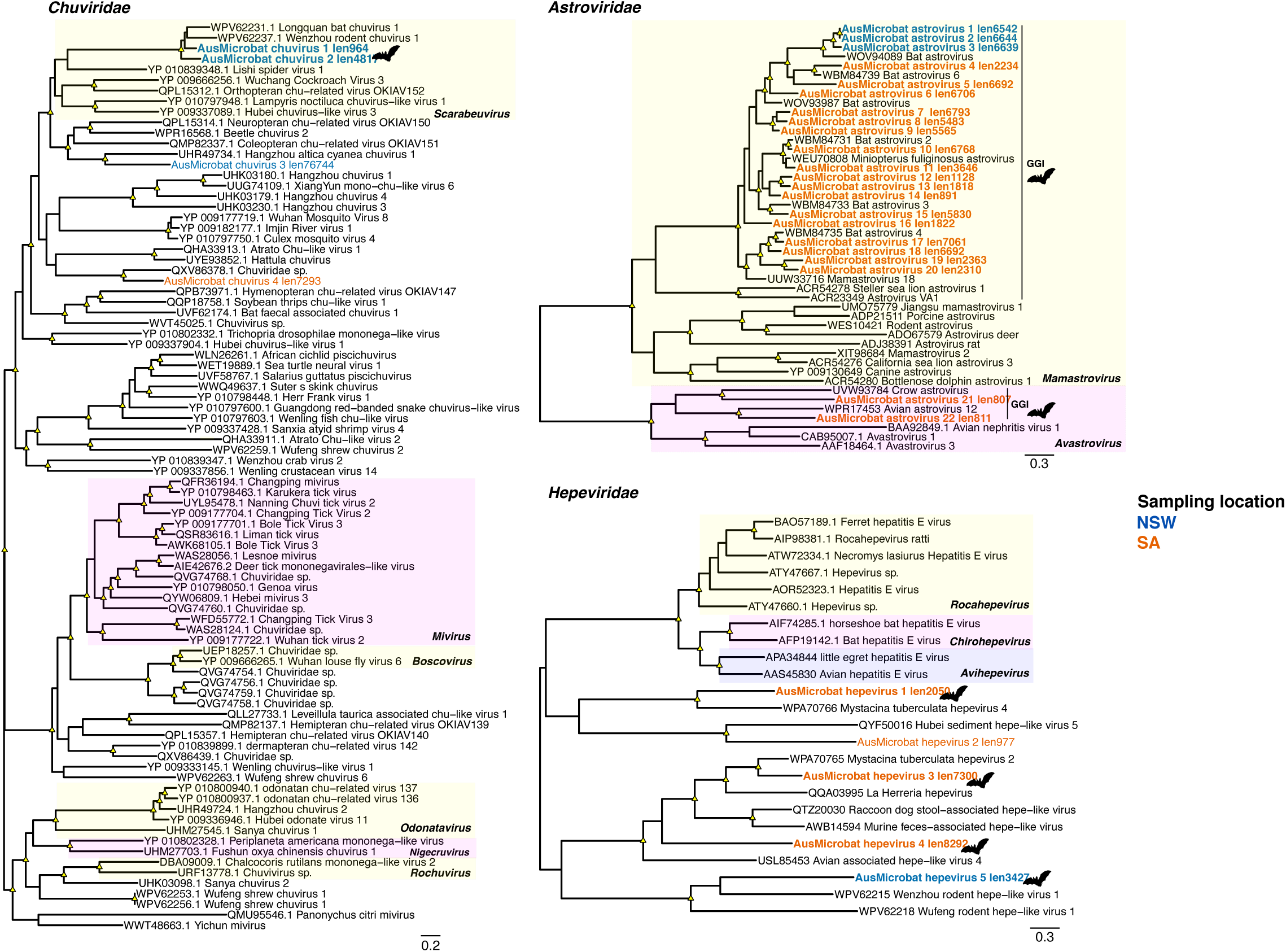
Phylogenetic relationships among the (A) *Chuviridae*, (B) *Astroviridae,* and (C) *Hepeviridae.* Trees are based on the amino acid sequences of the putative RdRp (*Chuviridae* and *Hepeviridae*) or capsid (*Astroviridae*). All phylogenetic trees are mid-point rooted for clarity only. Scale bars represent the number of amino acid substitutions per site. Known genera are represented by colour-coded clades for interpretation. Tip labels for novel viruses are coloured by sampling location – blue (NSW), orange (SA). Nodal support values ≥80% SH-aLRT and ≥95 % UFboot are denoted with yellow triangles at nodes. Tip labels highlighted in bold represent likely mammalian-associated lineages, also indicated with bat silhouettes.

**Table 2.** RNA viruses discovered in bat guano collected from sampling locations in NSW and SA, Australia. Taxonomic virus groups are indicated as well as the closest hit in the NCBI non-redundant (nr) database.

| **Virus** | **Family** | **Length** | **Accession code** | **Best hit (NCBI-nr)** | **% ID** | **E-value** | **Likely host** |
| --- | --- | --- | --- | --- | --- | --- | --- |
| AusMicrobat AlphaCov1 | *Coronaviridae* | 27784 | QGX41962.1 | polyprotein 1ab [Alphacoronavirus sp] | 81.0 | 0.0 | Mammal |
| AusMicrobat AlphaCov2 | *Coronaviridae* | 27366 | YP_010799723.1 | polyprotein 1ab [Alphacoronavirus sp] | 97.0 | 0.0 | Mammal |
| AusMicrobat AlphaCov3 | *Coronaviridae* | 27481 | QGX41962.1 | polyprotein 1ab [Alphacoronavirus sp] | 81.0 | 0.0 | Mammal |
| AusMicrobat AlphaCov4 | *Coronaviridae* | 2651 | YP_010037467.1 | polyprotein 1ab [Alphacoronavirus sp] | 99.5 | 0.0 | Mammal |
| AusMicrobat AlphaCov5 | *Coronaviridae* | 28670 | YP_010037467.1 | polyprotein 1ab [Alphacoronavirus sp] | 96.9 | 0.0 | Mammal |
| AusMicrobat AlphaCov6 | *Coronaviridae* | 29069 | WCZ55891.1 | MAG: ORF1a protein [Miniopterus bat coronavirus HKU8] | 73.4 | 0.0 | Mammal |
| AusMicrobat chuvirus 1 | *Chuviridae* | 964 | WPV62231.1 | MAG: RNA-dependent RNA polymerase, partial [Longquan bat chuvirus 1] | 80.9 | 72e-180 | Mammal |
| AusMicrobat chuvirus 2 | *Chuviridae* | 4817 | WNV56441.1 | MAG: RNA-dependent RNA polymerase [Wenzhou rodent chuvirus 1] | 65.3 | 0.0 | Mammal |
| AusMicrobat chuvirus 3 | *Chuviridae* | 7674 | UHR49734.1 | MAG: RNA-dependent RNA polymerase [Hangzhou altica cyanea chuvirus 1] | 42.6 | 0.0 | Invertebrate |
| AusMicrobat chuvirus 4 | *Chuviridae* | 7293 | WPR16568.1 | MAG: RNA-dependent RNA polymerase, partial [Beetle chuvirus 2] | 37.6 | 0.0 | Invertebrate |
| AusMicrobat astrovirus 1 | *Astroviridae* | 6542 | WNK76727.1 | MAG: capsid precursor protein [Bat astrovirus] | 61.0 | 61e-307 | Mammal |
| AusMicrobat astrovirus 2 | *Astroviridae* | 6644 | WNK76727.1 | MAG: capsid precursor protein [Bat astrovirus] | 61.0 | 4.98e-307 | Mammal |
| AusMicrobat astrovirus 3 | *Astroviridae* | 6639 | WNK76727.1 | MAG: capsid precursor protein [Bat astrovirus] | 61.0 | 5.66e-309 | Mammal |
| AusMicrobat astrovirus 4 | *Astroviridae* | 2234 | WNK76727.1 | MAG: capsid precursor protein [Bat astrovirus] | 58.8 | 6.91e-252 | Mammal |
| AusMicrobat astrovirus 5 | *Astroviridae* | 6692 | WBM84744.1 | MAG: capsid protein, partial [Bat astrovirus 10] | 84.7 | 0.0 | Mammal |
| AusMicrobat<br>astrovirus 6 | *Astroviridae* | 6706 | XBH23922.1 | MAG: capsid protein [Miniopterus bat<br>astrovirus] | 61.8 | 0.0 | Mammal |
| AusMicrobat<br>astrovirus 7 | *Astroviridae* | 6793 | XBH23922.1 | MAG: capsid protein [Miniopterus bat<br>astrovirus] | 75.3 | 0.0 | Mammal |
| AusMicrobat<br>astrovirus 8 | *Astroviridae* | 5483 | XBH23922.1 | MAG: capsid protein [Miniopterus bat<br>astrovirus] | 72.3 | 0.0 | Mammal |
| AusMicrobat<br>astrovirus 9 | *Astroviridae* | 5565 | XBH23922.1 | MAG: capsid protein [Miniopterus bat<br>astrovirus] | 74.0 | 0.0 | Mammal |
| AusMicrobat<br>astrovirus 10 | *Astroviridae* | 6768 | WBM84731.1 | MAG: capsid protein [Bat astrovirus 2] | 92.9 | 0.0 | Mammal |
| AusMicrobat<br>astrovirus 11 | *Astroviridae* | 3646 | WBM84731.1 | MAG: capsid protein [Bat astrovirus 2] | 85.3 | 0.0 | Mammal |
| AusMicrobat<br>astrovirus 12 | *Astroviridae* | 1128 | WEU70808.1 | MAG: capsid protein precursor, partial<br>[Miniopterus fuliginosus astrovirus] | 84.0 | 8.16e-210 | Mammal |
| AusMicrobat<br>astrovirus 13 | *Astroviridae* | 1818 | WBM84733.1 | MAG: capsid protein [Bat astrovirus 3] | 75.0 | 5.59e-279 | Mammal |
| AusMicrobat<br>astrovirus 14 | *Astroviridae* | 891 | WEU70808.1 | MAG: capsid protein precursor, partial<br>[Miniopterus fuliginosus astrovirus] | 86.6 | 3.25e-173 | Mammal |
| AusMicrobat<br>astrovirus 15 | *Astroviridae* | 5830 | WBM84733.1 | MAG: capsid protein [Bat astrovirus 3] | 75.1 | 0.0 | Mammal |
| AusMicrobat<br>astrovirus 16 | *Astroviridae* | 1822 | UUW33716.1 | ORF2, partial [Mamastrovirus 18] | 69.7 | 3.83e-289 | Mammal |
| AusMicrobat<br>astrovirus 17 | *Astroviridae* | 7061 | WBM84735.1 | MAG: capsid protein [Bat astrovirus 4] | 87.9 | 0.0 | Mammal |
| AusMicrobat<br>astrovirus 18 | *Astroviridae* | 6692 | WBM84735.1 | MAG: capsid protein [Bat astrovirus 4] | 82.0 | 0.0 | Mammal |
| AusMicrobat<br>astrovirus 19 | *Astroviridae* | 2363 | WBM84735.1 | MAG: capsid protein [Bat astrovirus 4] | 67.6 | 0.0 | Mammal |
| AusMicrobat<br>astrovirus 20 | *Astroviridae* | 2310 | WBM84735.1 | MAG: capsid protein [Bat astrovirus 4] | 69.6 | 0.0 | Mammal |
| AusMicrobat<br>astrovirus 21 | *Astroviridae* | 807 | UVW93784.1 | capsid protein, partial [Crow astrovirus] | 52.6 | 3.75e-68 | Mammal |
| AusMicrobat<br>astrovirus 22 | *Astroviridae* | 811 | WPR17453.1 | MAG: capsid protein [Avian astrovirus 12] | 52.8 | 2.64e-85 | Mammal |
| AusMicrobat hepevirus 1 | *Hepeviridae* | 2050 | WPA70766.1 | MAG: nonstructural polyprotein, partial [Mystacina tuberculata hepevirus 4] | 45.2 | 6.86e-179 | Mammal |
| AusMicrobat hepevirus 3 | *Hepeviridae* | 7300 | QQA03995.1 | polyprotein [La Herreria hepevirus] | 41.3 | 0.0 | Mammal |
| AusMicrobat hepevirus 4 | *Hepeviridae* | 8292 | QQA03995.1 | polyprotein [La Herreria hepevirus] | 30.8 | 2.52e-195 | Mammal |
| AusMicrobat hepevirus 5 | *Hepeviridae* | 3427 | WPV62215.1 | WPV62215.1 MAG: non-structural polyprotein, partial [Wenzhou rodent hepe-like virus 1] | 34.0 | 4.87e-103 | Mammal |
| AusMicrobat hepevirus 2 | *Hepeviridae* | 977 | QYF50016.1 | MAG: replicase, partial [Hubei sediment hepe-like virus 5] | 37.2 | 1.44e-50 | Invertebrate |
| AusMicrobat nodavirus 7 | *Nodaviridae* | 2707 | WPA70719.1 | MAG: RNA-dependent RNA polymerase, partial [Mystacina tuberculata nodavirus 4] | 42.5 | 1.37e-91 | Mammal |
| AusMicrobat nodavirus 1 | *Nodaviridae* | 3265 | QBS55246.1 | RNA-dependent RNA polymerase [Tetranychus urticae-associated nodavirus A] | 62.0 | 0.0 | Invertebrate |
| AusMicrobat nodavirus 2 | *Nodaviridae* | 3410 | QUS52660.1 | RNA-dependent RNA polymerase [Mute swan feces associated noda-like virus 2] | 67.0 | 0.0 | Invertebrate |
| AusMicrobat nodavirus 3 | *Nodaviridae* | 8454 | WPV63042.1 | MAG: RNA-dependent RNA polymerase [Wufeng bat nodavirus 1] | 34.2 | 1.86e-138 | Invertebrate |
| AusMicrobat nodavirus 4 | *Nodaviridae* | 4667 | ADI48250.1 | putative RdRp [Bat guano associated nodavirus GF-4n] | 43.1 | 1.93e-215 | Invertebrate |
| AusMicrobat nodavirus 5 | *Nodaviridae* | 2812 | ASU47554.1 | polymerase [Lone star tick nodavirus] | 41.2 | 3.71e-173 | Invertebrate |
| AusMicrobat nodavirus 6 | *Nodaviridae* | 3763 | UQB76069.1 | RNA-dependent RNA polymerase [Flumine noda-like virus 3] | 39.1 | 5.30e-186 | Invertebrate |
| AusMicrobat nodavirus 8 | *Nodaviridae* | 4993 | XHA85947.1 | MAG: putative RNA-dependent RNA polymerase [Qianjiang noda-like virus 29] | 36.5 | 3.98e-95 | Invertebrate |
| AusMicrobat nodavirus 9 | *Nodaviridae* | 5407 | XHA85922.1 | MAG: hypothetical protein [Qianjiang noda-like virus 20] | 98.3 | 0.0 | Invertebrate |
| AusMicrobat nodavirus 10 | *Nodaviridae* | 6478 | WPV63071.1 | MAG: RNA-dependent RNA polymerase [Wufeng shrew nodamuvirus 1] | 38.4 | 9.20e-103 | Invertebrate |
| AusMicrobat nodavirus 11 | *Nodaviridae* | 3092 | WAX26160.1 | MAG: RNA-dependent RNA polymerase [Army ant associated Nodavirus 1] | 70.7 | 0.0 | Invertebrate |
| AusMicrobat nodavirus 12 | *Nodaviridae* | 5532 | WPA70725.1 | MAG: RNA-dependent RNA polymerase [Bat nodavirus] | 93.5 | 0.0 | Invertebrate |
| AusMicrobat<br>picornavirus 9 | *Picornavirales* | 5366 | WPV63438.1 | MAG: RNA-dependent RNA polymerase<br>[Longquan rodent picorna-like virus 1] | 54.3 | 0.0 | Mammal |
| AusMicrobat<br>picornavirus 10 | *Picornavirales* | 11453 | WFG77351.1 | MAG: polyprotein [Bat picornavirus 5] | 69.5 | 0.0 | Mammal |
| AusMicrobat<br>picornavirus 15 | *Picornavirales* | 7646 | AEM23660.1 | polyprotein [Bat picornavirus 2] | 78.4 | 0.0 | Mammal |
| AusMicrobat<br>picornavirus 16 | *Picornavirales* | 7727 | YP_004782528.1 | polyprotein [Bat picornavirus 1] | 79.1 | 0.0 | Mammal |
| AusMicrobat<br>picornavirus 1 | *Picornavirales* | 11568 | UDL13970.1 | MAG: polyprotein [Xiangshan picorna-like virus<br>4] | 30.2 | 0.0 | Invertebrate |
| AusMicrobat<br>picornavirus 2 | *Picornavirales* | 15044 | YP_009337161.1 | hypothetical protein [Hubei picorna-like virus<br>27] | 39.5 | 0.0 | Invertebrate |
| AusMicrobat<br>picornavirus 3 | *Picornavirales* | 10580 | YP_009110667.1 | polyprotein [Laodelphax striatellus picorna-like<br>virus 2] | 66.8 | 0.0 | Invertebrate |
| AusMicrobat<br>picornavirus 4 | *Picornavirales* | 11286 | UDL13967.1 | MAG: polyprotein [Xiangshan picorna-like virus<br>1] | 30.5 | 0.0 | Invertebrate |
| AusMicrobat<br>picornavirus 5 | *Picornavirales* | 1319 | WIW43234.1 | MAG: putative polyprotein, partial [Pteropus<br>rufus picorna-like virus] | 92.9 | 1.22e-225 | Invertebrate |
| AusMicrobat<br>picornavirus 6 | *Picornavirales* | 6473 | QKK82954.1 | hypothetical protein, partial [Erigeron annuus<br>picorna-like virus] | 90.9 | 0.0 | Invertebrate |
| AusMicrobat<br>picornavirus 7 | *Picornavirales* | 9422 | QZZ63320.1 | hypothetical protein [Nelson Picorna-like virus<br>3] | 74.5 | 0.0 | Invertebrate |
| AusMicrobat<br>picornavirus 8 | *Picornavirales* | 9857 | UUV42601.1 | MAG: polyprotein [Tuatara cloaca-associated<br>picorna-like virus-1] | 32.5 | 1.65e-269 | Invertebrate |
| AusMicrobat<br>picornavirus 11 | *Picornavirales* | 11017 | WPR16530.1 | MAG: RNA-dependent RNA polymerase, partial<br>[Avian associated picorna-like virus 41] | 49.3 | 0.0 | Invertebrate |
| AusMicrobat<br>picornavirus 12 | *Picornavirales* | 1905 | UYL83183.1 | MAG: RNA-dependent RNA polymerase, partial<br>[XiangYun tombus-noda-like virus 5] | 51.6 | 1.67e-21 | Invertebrate |
| AusMicrobat<br>picornavirus 13 | *Picornavirales* | 2727 | QUS52703.1 | polyprotein [Mute swan feces associated<br>picorna-like virus 7] | 31.0 | 5.15e-94 | Invertebrate |
| AusMicrobat<br>picornavirus 14 | *Picornavirales* | 6533 | QED42981.1 | hypothetical protein, partial [Pleurotus<br>picornavirus A] | 44.4 | 0.0 | Invertebrate |
| AusMicrobat<br>picornavirus 17 | *Picornavirales* | 11194 | WBM81658.1 | MAG: RNA-dependent RNA polymerase<br>[Myotis brandtii picorna-like virus 1] | 31.7 | 2.78e-212 | Invertebrate |
| AusMicrobat<br>picornavirus 18 | *Picornavirales* | 6502 | WPV63421.1 | MAG: RNA-dependent RNA polymerase<br>[Jingmen shrew picorna-like virus 1] | 30.4 | 1.29e-284 | Invertebrate |
| AusMicrobat<br>picornavirus 19 | *Picornavirales* | 9276 | UFZ21066.1 | MAG: structural polyprotein [Planococcus ficus-<br>associated picorna-like virus 1] | 46.2 | 5.82e-209 | Invertebrate |
| AusMicrobat<br>picornavirus 20 | *Picornavirales* | 7967 | WPV63450.1 | MAG: RNA-dependent RNA polymerase, partial<br>[Wenzhou bat picorna-like virus 8] | 43.1 | 2.20e-56 | Invertebrate |
| AusMicrobat<br>picornavirus 21 | *Picornavirales* | 11716 | UXD80005.1 | putative non-structural polyprotein [Lasius<br>neglectus picorna-like virus 3] | 46.6 | 5.11e-286 | Invertebrate |
| AusMicrobat<br>picornavirus 22 | *Picornavirales* | 10855 | YP_009336820.1 | hypothetical protein [Hubei picorna-like virus<br>53] | 30.0 | 0.0 | Invertebrate |
| AusMicrobat<br>picornavirus 23 | *Picornavirales* | 12484 | YP_009337118.1 | hypothetical protein [Hubei picorna-like virus<br>52] | 33.8 | 0.0 | Invertebrate |
| AusMicrobat<br>picornavirus 24 | *Picornavirales* | 12831 | UDL13977.1 | MAG: RNA dependent RNA polymerase<br>[Xiangshan picorna-like virus 7] | 25.2 | 4.24e-130 | Invertebrate |
| AusMicrobat<br>picornavirus 25 | *Picornavirales* | 1997 | AQY59901.1 | RdRp, partial [Statovirus B1] | 28.2 | 8.48e-46 | Invertebrate |
| AusMicrobat<br>picornavirus 26 | *Picornavirales* | 11186 | UQZ09572.1 | putative polyprotein [Freshwater macrophyte<br>associated picorna-like virus 12] | 25.3 | 1.71e-98 | Invertebrate |
| AusMicrobat<br>picornavirus 27 | *Picornavirales* | 6340 | XEQ84329.1 | hypothetical protein [Mink picorna-like virus 1] | 34.3 | 2.86e-123 | Invertebrate |
| AusMicrobat<br>picornavirus 28 | *Picornavirales* | 5594 | UXD80013.1 | putative non-structural polyprotein [Myrmica<br>rubra picorna-like virus 4] | 52.6 | 0.0 | Invertebrate |
| AusMicrobat<br>calicivirus 1 | *Picornavirales:*<br>*Caliciviridae* | 6514 | WAP91247.1 | MAG: nonstructural polyprotein, partial [Avian<br>associated calicivirus 5] | 46.8 | 0.0 | Mammal |
| AusMicrobat<br>calicivirus 2 | *Picornavirales:*<br>*Caliciviridae* | 10043 | WAP91247.1 | MAG: nonstructural polyprotein, partial [Avian<br>associated calicivirus 5] | 46.7 | 0.0 | Mammal |
| AusMicrobat<br>calicivirus 3 | *Picornavirales:*<br>*Caliciviridae* | 10146 | USL85468.1 | MAG: non-structural polyprotein [Avian<br>associated calicivirus 1] | 39.2 | 0.0 | Mammal |
| AusMicrobat<br>rhabdovirus 3 | *Rhabdoviridae* | 1205 | WWV88395.1 | MAG: L protein, partial [Rhabdovirus sp.] | 71.6 | 5.49e-201 | Mammal |
| AusMicrobat<br>rhabdovirus 4 | *Rhabdoviridae* | 2488 | WWV88387.1 | MAG: L protein [Rhabdovirus sp.] | 65.9 | 0.0 | Mammal |
| AusMicrobat<br>rhabdovirus 12 | *Rhabdoviridae* | 3510 | WPV62811.1 | MAG: RNA-dependent RNA polymerase<br>[Wenzhou bat rhabdovirus 2] | 72.2 | 0.0 | Mammal |
| AusMicrobat rhabdovirus 13 | *Rhabdoviridae* | 2288 | WPV62809.1 | MAG: putative glycoprotein [Wenzhou bat rhabdovirus 2] | 68.4 | 3.88e-113 | Mammal |
| AusMicrobat rhabdovirus 1 | *Rhabdoviridae* | 770 | WPV62848.1 | MAG: RNA-dependent RNA polymerase, partial [Wufeng shrew rhabdovirus 15] | 59.4 | 5.49e-90 | Invertebrate |
| AusMicrobat rhabdovirus 2 | *Rhabdoviridae* | 764 | QQP18754.1 | RNA-dependent RNA polymerase [Soybean thrips rhabdo-like virus 2] | 52.7 | 2.70e-82 | Invertebrate |
| AusMicrobat rhabdovirus 5 | *Rhabdoviridae* | 13603 | YP_009204560.1 | L protein [Fox fecal rhabdovirus] | 33.5 | 8.48e-256 | Invertebrate |
| AusMicrobat rhabdovirus 6 | *Rhabdoviridae* | 1415 | QMP82309.1 | RNA-dependent RNA polymerase, partial [Hemipteran rhabdo-related virus OKIAV26] | 57.6 | 5.05e-169 | Invertebrate |
| AusMicrobat rhabdovirus 7 | *Rhabdoviridae* | 14451 | QMP82194.1 | RNA-dependent RNA polymerase [Coleopteran rhabdo-related virus OKIAV28] | 34.0 | 0.0 | Invertebrate |
| AusMicrobat rhabdovirus 8 | *Rhabdoviridae* | 13467 | WGF21055.1 | L [Spodoptera frugiperda rhabdovirus]<br>[Spodoptera frugiperda rhabdovirus] | 82.9 | 0.0 | Invertebrate |
| AusMicrobat rhabdovirus 9 | *Rhabdoviridae* | 9821 | YP_010798566.1 | RNA-dependent RNA polymerase [Lepidopteran rhabdo-related virus 34] | 47.4 | 0.0 | Invertebrate |
| AusMicrobat rhabdovirus 10 | *Rhabdoviridae* | 1040 | DAZ87777.1 | TPA_asm: hypothetical protein [Metorhabdovirus 1] | 78.0 | 3.63e-180 | Invertebrate |
| AusMicrobat rhabdovirus 11 | *Rhabdoviridae* | 1480 | DAZ87842.1 | TPA_asm: hypothetical protein [Clonorhabdovirus 1] | 77.9 | 7.18e-263 | Invertebrate |
| AusMicrobat phenuivirus 1 | *Bunyavirales* | 1976 | WPV62188.1 | MAG: RNA-dependent RNA polymerase [Wufeng shrew phenuivirus 9] | 78.1 | 0.0 | Mammal |
| AusMicrobat phenuivirus 2 | *Bunyavirales* | 1140 | WPV62186.1 | MAG: RNA-dependent RNA polymerase [Wufeng shrew phenuivirus 7] | 69.4 | 1.19e-173 | Mammal |
| AusMicrobat phenuivirus 10 | *Bunyavirales* | 741 | WPV62168.1 | MAG: RNA-dependent RNA polymerase [Wenzhou bat phenuivirus 2] | 60.3 | 4.24e-94 | Mammal |
| AusMicrobat phenuivirus 3 | *Bunyavirales* | 702 | DBA56514.1 | TPA_asm: RNA-dependent RNA polymerase [Neotermes castaneus phenuivirus 1] | 51.3 | 1.06e-63 | Invertebrate |
| AusMicrobat phenuivirus 4 | *Bunyavirales* | 850 | QOR29577.1 | RNA-dependent RNA polymerase, partial [Bat bunyavirus] | 57.9 | 1.03e-99 | Invertebrate |
| AusMicrobat phenuivirus 5 | *Bunyavirales* | 1541 | APL98134.1 | RNA-dependent RNA polymerase, partial [Bat bunyavirus JTM] | 56.2 | 1.47e-174 | Invertebrate |
| AusMicrobat phenuivirus 6 | *Bunyavirales* | 7039 | UVZ34188.1 | RNA-dependent RNA polymerase [Erthesina fullo bunyavirus 1] | 39.5 | 0.0 | Invertebrate |
| AusMicrobat<br>phenuivirus 7 | *Bunyavirales* | 7570 | QMP82296.1 | RNA-dependent RNA polymerase [Lepidopteran<br>phenui-related virus OKIAV270] | 58.4 | 0.0 | Invertebrate |
| AusMicrobat<br>phenuivirus 8 | *Bunyavirales* | 9004 | XGY11997.1 | MAG: RNA-dependent RNA polymerase<br>[Empoasca fabae phenuivirus 1] | 42.5 | 1.90e-212 | Invertebrate |
| AusMicrobat<br>phenuivirus 9 | *Bunyavirales* | 6996 | QMP82311.1 | RNA-dependent RNA polymerase [Blattodean<br>phenui-related virus OKIAV261] | 40.9 | 0.0 | Invertebrate |
| AusMicrobat<br>phenuivirus 11 | *Bunyavirales* | 989 | XCN99621.1 | RNA-dependent RNA polymerase [Switchgrass<br>phenui-like virus 1] | 59.6 | 6.01e-130 | Invertebrate |
| AusMicrobat<br>bunyavirus 1 | *Bunyavirales* | 794 | WPV62125.1 | MAG: RNA-dependent RNA polymerase<br>[Jingmen bat bunyavirus 1] | 75.8 | 1.10e-126 | Mammalian |
| AusMicrobat<br>bunyavirus 2 | *Bunyavirales* | 2868 | AOA33722.1 | putative RNA dependent RNA polymerase<br>[Crithidia otongatchiensis leishbunyavirus<br>CotoLBV1] | 49.8 | 4.92e-289 | Invertebrate |
| AusMicrobat<br>bunyavirus 3 | *Bunyavirales* | 1881 | UPT53676.1 | MAG: RNA-dependent RNA polymerase<br>[Bactrocera correcta trypanosomatid<br>leishbunyavirus] | 56.5 | 4.98e-234 | Invertebrate |
| AusMicrobat<br>bunyavirus 4 | *Bunyavirales* | 1439 | APG79301.1 | RNA-dependent RNA polymerase, partial<br>[Hubei bunya-like virus 5] | 45.3 | 8.94e-124 | Invertebrate |
| AusMicrobat<br>bunyavirus 5 | *Bunyavirales* | 7763 | DBA56512.1 | TPA_asm: RNA-dependent RNA polymerase<br>[Neotermes castaneus bunya-like virus 1] | 39.2 | 0.0 | Invertebrate |
| AusMicrobat<br>sedoreovirus 1 | *Sedoreoviridae* | 4013 | WPV63639.1 | MAG: RNA-dependent RNA polymerase, partial<br>[Jingmen bat reo-like virus 1] | 36.40 | 1.48e-241 | Mammal |
| AusMicrobat<br>sedoreovirus 2 | *Sedoreoviridae* | 3991 | APG79114.1 | RdRp [Hubei reo-like virus 12] | 36.70 | 2.08e-223 | Mammal |
| AusMicrobat<br>sedoreovirus 3 | *Sedoreoviridae* | 2031 | AVM87459.1 | RNA-dependent RNA polymerase [Wenling<br>scaldfish reovirus] | 26.20 | 2.04e-57 | Mammal |
| AusMicrobat<br>sedoreovirus 4 | *Sedoreoviridae* | 2655 | XBH23973.1 | MAG: VP1 protein, partial [Mops bat reovirus] | 46.70 | 3.11e-272 | Mammal |
| AusMicrobat<br>sedoreovirus 5 | *Sedoreoviridae* | 636 | QCX36715.1 | structural protein VP1, partial [Human<br>rotavirus] | 99.10 | 3.27e-139 | Mammal |
| AusMicrobat<br>sedoreovirus 6 | *Sedoreoviridae* | 3640 | UUW33709.1 | VP1 protein [Rotavirus J] | 92.70 | 0.0 | Mammal |
| AusMicrobat<br>sedoreovirus 7 | *Sedoreoviridae* | 731 | AWV67027.1 | VP1, partial [Bat rotavirus H] | 75.60 | 7.90e-116 | Mammal |
| AusMicrobat<br>sedoreovirus 7 | *Sedoreoviridae* | 1979 | AWV67027.1 | VP1, partial [Bat rotavirus H] | 82.10 | 0.0 | Mammal |
| AusMicrobat<br>sedoreovirus 7 | *Sedoreoviridae* | 6118 | AWV67027.1 | VP1, partial [Bat rotavirus H] | 81.00 | 0.0 | Mammal |
| AusMicrobat<br>sedoreovirus 7 | *Sedoreoviridae* | 5485 | AWV67027.1 | VP1, partial [Bat rotavirus H] | 81.10 | 0.0 | Mammal |
| AusMicrobat<br>sedoreovirus 7 | *Sedoreoviridae* | 11121 | AWV67027.1 | VP1, partial [Bat rotavirus H] | 81.00 | 0.0 | Mammal |
| AusMicrobat<br>sedoreovirus 8 | *Sedoreoviridae* | 4253 | WPV63640.1 | MAG: RNA-dependent RNA polymerase<br>[Jingmen bat reo-like virus 2] | 23.10 | 9.08e-55 | Invertebrate |
| AusMicrobat<br>sedoreovirus 8 | *Sedoreoviridae* | 4254 | WPV63640.1 | MAG: RNA-dependent RNA polymerase<br>[Jingmen bat reo-like virus 2] | 23.10 | 9.10e-55 | Invertebrate |
| AusMicrobat<br>sedoreovirus 9 | *Sedoreoviridae* | 4258 | UOI84724.1 | RNA-dependent RNA polymerase [Ceratitis<br>capitata reo-like virus 1] | 33.10 | 3.78e-188 | Invertebrate |
| AusMicrobat<br>sedoreovirus 10 | *Sedoreoviridae* | 1608 | APG79196.1 | RNA-dependent RNA polymerase [Hubei reo-like virus 6] | 34.70 | 2.13e-83 | Invertebrate |
| AusMicrobat<br>sedoreovirus 11 | *Sedoreoviridae* | 2000 | APG79200.1 | RdRp [Hubei reo-like virus 5] | 23.70 | 3.49e-23 | Invertebrate |
| AusMicrobat<br>sedoreovirus 12 | *Sedoreoviridae* | 1930 | APG79166.1 | RdRp [Hubei reo-like virus 9] | 25.40 | 8.70e-29 | Invertebrate |

To investigate the potential co-circulation of viruses at a single location, we focused on the virus profile found in bat faecal samples collected from the Naracoorte bat maternity caves as it houses a single bat species. Accordingly, we detected diverse mammalian-associated viruses co-circulating in the Southern bent-wing bat population, including a high diversity of mamastroviruses, but also rhabdoviruses, bunyaviruses, coronaviruses, hepeviruses, nodaviruses, picornaviruses and reoviruses shed in bat guano (**Figure 2-4**, **Figure 5**).

**Figure 4.**
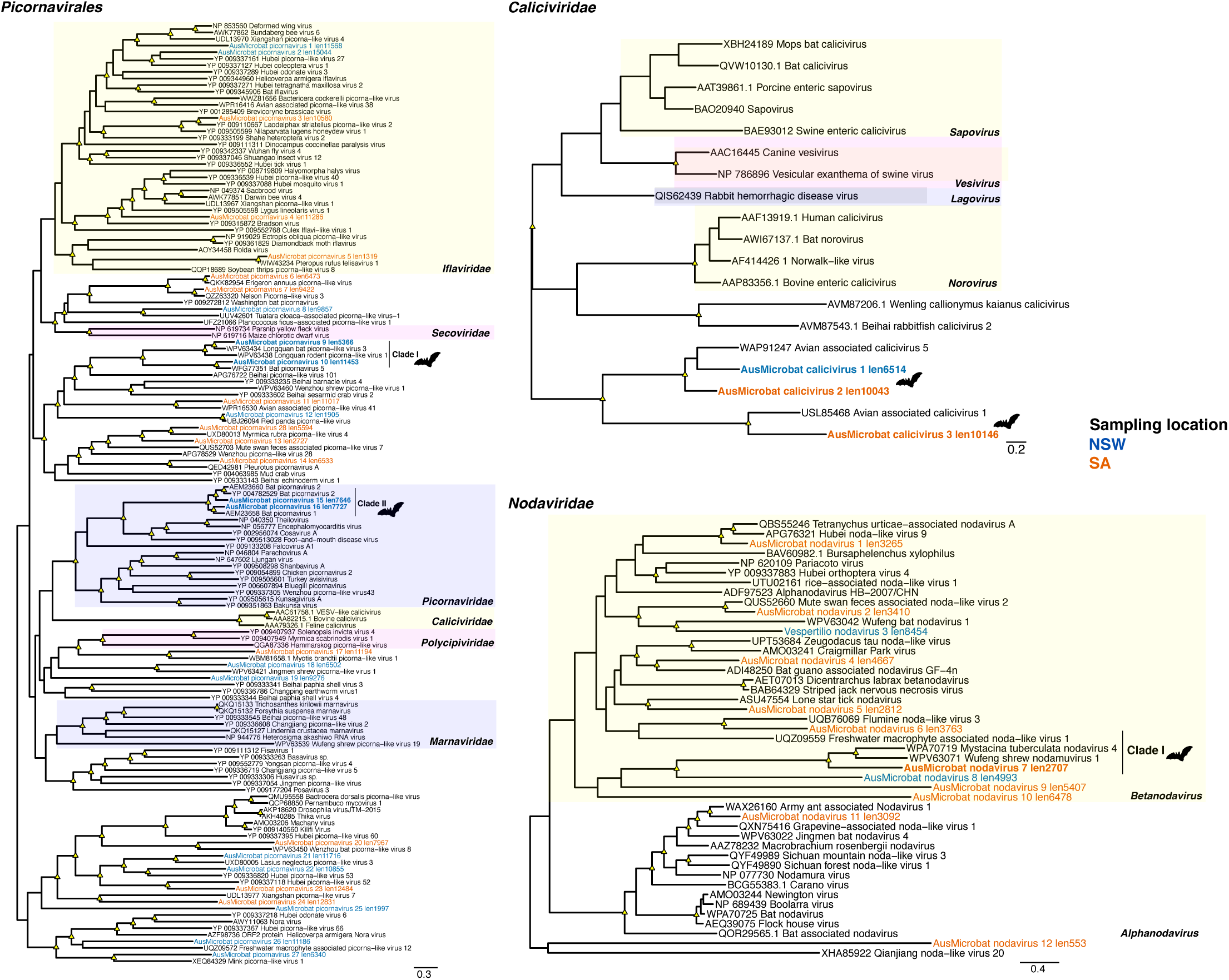
Phylogenetic relationships among the (A) *Picornavirales* (B) *Caliciviridae*, and (C) *Nodaviridae*. Trees are based on the amino acid sequences of the putative RdRp. All phylogenetic trees are mid-point rooted for clarity only. Scale bars represent the number of amino acid substitutions per site. Known genera are represented by colour-coded clades for easier interpretation. Tip labels for novel viruses are coloured by sampling location – blue (NSW), orange (SA). Nodal support values ≥80% SH-aLRT and ≥95 % UFboot are denoted with yellow triangles at nodes. Tip labels highlighted in bold represent likely mammalian-associated lineages, also indicated with bat silhouettes.

**Figure 5.**
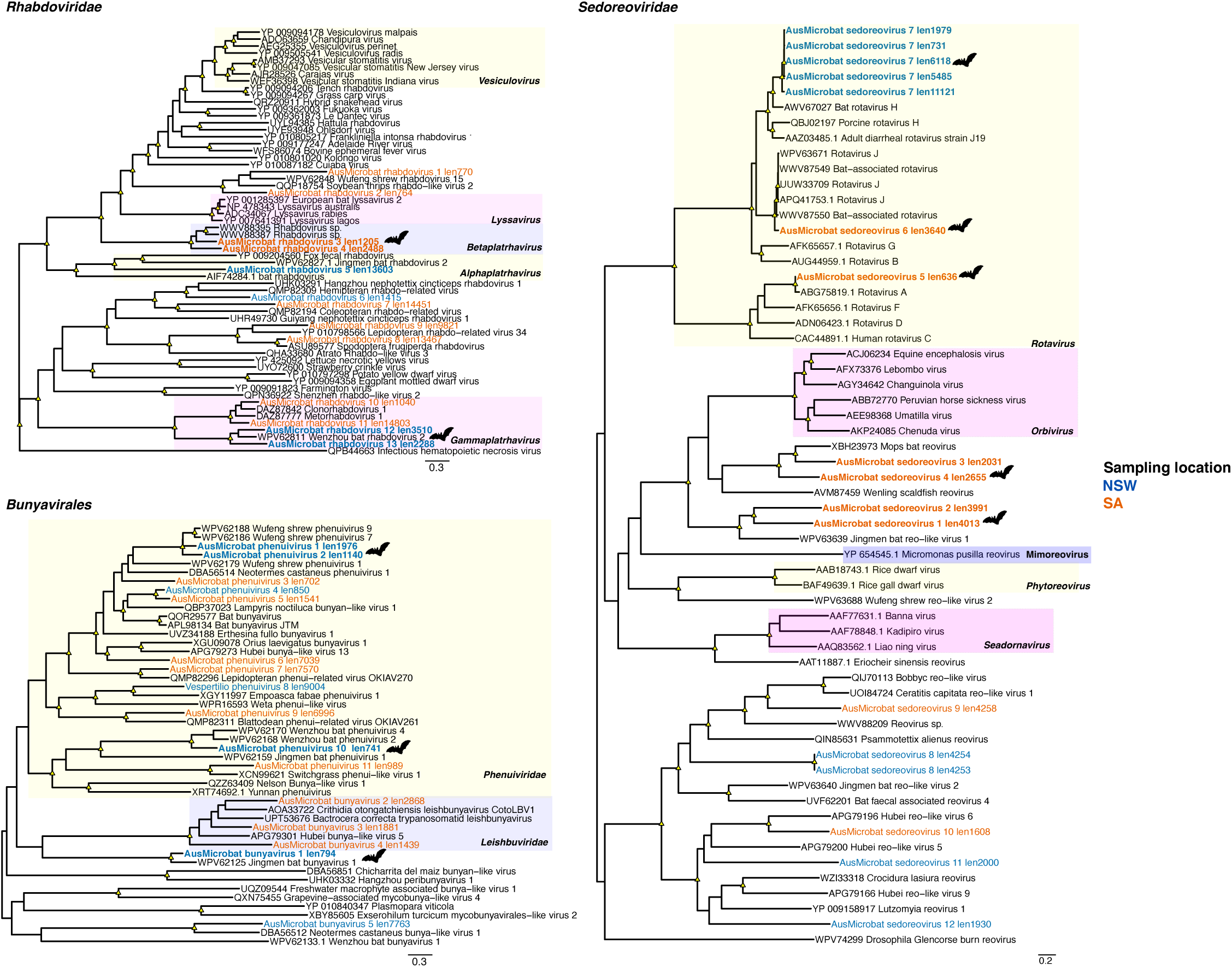
Phylogenetic relationships among the (A) *Bunyavirales* and (B) *Sedoreoviridae*. Trees are based on the amino acid sequences of the putative RdRp. All phylogenetic trees are mid-point rooted for clarity only. Scale bars represent the number of amino acid substitutions per site. Known genera are represented by colour-coded clades for easier interpretation. Tip labels for novel viruses are coloured by sampling location – blue (NSW), orange (SA). Nodal support values ≥80% SH-aLRT and ≥95 % UFboot are denoted with yellow triangles at nodes. Tip labels highlighted in bold represent likely mammalian-associated lineages, also indicated with bat silhouettes.

### RNA virus families

#### Coronaviridae

A striking result of this study was the relatively large number (six) of alphacoronaviruses identified in microbat faecal samples from NSW and SA. In marked contrast, no betacoronaviruses were identified in our sampling. The bat alphacoronaviruses identified here fell into three subgenera: *Nyctacovirus*, *Minunacovirus*, and *Pedacovirus* (**Figure 2A-B**). As expected, the genome sequences of the newly identified alphacoronaviruses ranged from 27.36 – 32.02 Kb and exhibited the genomic configurations expected for these viruses. The coronaviruses detected in faecal samples from NSW were highly abundant (TPM = 13.2 – 418.43) in comparison to those found in SA samples (TPM = 0.23 – 3.4), perhaps reflecting local aspects of virus ecology or sampling artifacts. Of note, the newly identified members of the subgenus *Nyctacovirus* were closely related (97.2 – 99.77% amino acid identity in the RdRp) to those found in different microbat species sampled in Western Australian, Queensland, as well as Christmas Island, and hence indicative of a broad geographic distribution. Similarly, the alphacoronavirus sequences from the subgenus *Pedacovirus* were closely related to an alphacoronavirus identified in Western Australia (**Figure 2A-B**).

AusMicrobat alphacoronavirus 6 (TPM = 3.4) from the subgenus *Minunacovirus* was closely related to sequences circulating in Queensland and Southern Asia, including *Miniopterus* bat coronavirus HKU8 previously detected in bat faecal samples from Hong Kong (97.6% amino acid identity in the RdRp) and associated with rhinolopid bats (**Figure 2A-B**) [36]. These geographic patterns were consistent with the clustering analysis, in which clusters contained sequences from various locations across Australia, including NSW and SA (**Figure 2C**).

#### Chuviridae

Also of significance was the identification of four chuviruses (*Chuviridae*) in microbat faeces collected from bat boxes in NSW (n = 3) and SA (n = 1) and provisionally termed AusMicrobat chuviruses 1-4. These sequences shared up to 78% amino acid identity among themselves and grouped with viruses from the genus *Scarabeuvirus* as well as with unclassified chuviruses (TPM = 0.34 – 5.75) (**Figure 3A**). Of most note, in our phylogenetic analysis AusMicrobat chuvirus 1-2 (TPM = 0.34 – 0.52) clustered with Longquan bat chuvirus 1 and Wenzhou rodent chuvirus 1 (∼92 – 95% amino acid identity in the RdRp), previously sampled from the gut and lung of the Chinese rufous horseshoe bat (*Rhinolophus sinicus*) and the striped field mouse (*Apodemus agrarius*) [37], respectively. Together, these four sequences formed a distinct mammalian lineage within the *Chuviridae*. As such, this provides strong evidence for the sustained transmission of exogenous chuviruses in mammals on multiple continents. In contrast, AusMicrobat chuvirus 3 grouped with invertebrate-associated viruses primarily detected in coleopterans, while AusMicrobat chuvirus 4 formed a lineage with a viral sequence identified in an avian faecal metagenome, indicative of a dietary association.

#### Astroviridae

We identified a high diversity of astroviruses (n = 22) that clustered with bat viruses within the genus *Mamastrovirus* (TPM = 0.46 – 138), with genomes ranging from ∼5-7 kb in length. The majority of these viruses were detected in samples from the Naracoorte maternity caves in SA (91 – 98% amino acid identity) and were related to sequences found in intestine, skin, and swabs samples from *Miniopterus* spp. (*Miniopteridae*) from Australia (NSW) and China (59 – 93% amino acid identity). The remaining mamastroviruses formed a clade comprising sequences from NSW (TPM = 0.46 – 1.29), which grouped with a Chinese sequence detected in the intestine of a *Pipistrellus pipistrellus* bat. Finally, two other novel astroviruses were detected at low abundance levels and grouped within the genus *Avastrovirus* (TPM = 0.08 – 0.18) (**Figure 3B**), primarily associated with infection in birds (∼53% amino acid identity). Whether this reflects cross-species transmission between bats and birds is uncertain.

#### Hepeviridae

The majority of the newly discovered hepeviruses were related to those found in vertebrate metagenomes (**Figure 3C**), suggesting a mammalian origin. This was exemplified by AusMicrobat hepevirus 5 (TPM = 0.38) which grouped with rodent hepeviruses in our phylogenetic analysis. Notably, two of the novel hepeviruses from the SA samples were related to viruses found in bat guano from lesser short-tailed bats (*Mystacina tuberculata*) (TPM = 0.18 – 9.24), one of the only two native terrestrial mammals in New Zealand (31-63% amino acid identity). This could reflect either shared arthropod diets between Australian and New Zealand bats or virus-host co-divergence reflecting the separation of Australian and New Zealand bats some 16-18 million years ago [23, 38]. In contrast, AusMicrobat hepevirus 2 grouped with a virus previously identified in sediment in China (19% aa identity).

#### Picornavirales

We identified 30 viral contigs distributed among different families within the order *Picornavirales* (**Figure 4A**), including the *Picornaviridae*, *Caliciviridae*, *Marnaviridae*, *Polycipiviridae*, *Iflaviridae* as well as previously unclassified viruses. The novel viruses exhibited amino acid identities to their closest relatives ranging from 25–93% in the RdRp (**Table 2**). Phylogenetic analysis indicated that AusMicrobat picornaviruses 9,10,15 and 16 were likely associated with mammals, grouping with sequences identified in faeces from bats and rodents (TPM = 7.37 – 2199) (**Figure 4A**; Clade II-II), whereas the remaining virus sequences were grouped with invertebrate-associated viruses and hence are likely dietary associated. Similarly, we identified three caliciviruses from samples collected in NSW and SA that shared 17–52% amino acid sequence identity between them in the RdRp (**Figure 4B**). Only one calicivirus sequence was identified in a bat box from NSW (TPM = 0.8). However, the novel caliciviruses did not cluster with bat caliciviruses but instead formed a sister clade with avian caliciviruses (up to 39% RdRp amino acid identity) (**Figure 4B)**.

#### Nodaviridae

We identified a large number of nodavirus sequences (2707–8454 bp) from bat faecal samples that were related to invertebrate viruses within the genus *Alphanodavirus*, sharing between 36–70% amino acid sequence identity in the RdRp with their closest relatives (**Figure 4C**). However, AusMicrobat nodavirus 7 (TPM = 0.77) fell within a mammalian-related clade grouping with bat and shrew metagenome-derived sequences (Clade I), including viral sequences found in the New Zealand lesser short-tailed bat (*Mystacina tuberculata*) and the Asian gray shrew (*Crocidura attenuate*), indicating both shared common ancestry and a likely mammalian origin.

#### Rhabdoviridae

We detected 13 novel rhabdoviruses, including five likely bat-associated viruses related to mammalian-derived sequences from bats and shrews (Figure 5**A**). The newly identified AusMicrobat rhabdoviruses shared between 33–83% amino acid identity in the RdRp with their closest relatives, whereas their relative abundance levels ranged from 0.52–5.84 TPM (**Table 2**). In particular, the likely bat-associated viruses were related to metagenome-derived sequences previously identified from bats in China within the *Alpha*, *Beta* and *Gammaplatrhavirus* genera. For instance, AusMicrobat rhabdoviruses 3-4 (TPM = 1.08 – 1.26) formed a lineage with bat metagenome-derived sequences from the intestine and faecal samples of *Miniopterus pusillus* and *Hipposideros armiger* in China (66–72% amino acid identity in the RdRp), suggesting a broad circulation of rhabdoviruses in bats. Finally, invertebrate-associated rhabdoviruses were related to viral sequences detected in invertebrates as well as plants.

#### Bunyavirales

A total of 16 bunyaviruses were found in bat faecal samples collected from NSW and SA, with RdRp amino acid identities of 39-83% to already documented bunyavirus sequences (TPM = 0.11 – 760.29). Newly discovered bunyaviruses fell within diverse lineages from the families *Phenuiviridae* and *Leishbuviridae,* as well as a number of unclassified lineages (**Figure 5B**). AusMicrobat phenuiviruses 1, 2 and 10, as well as AusMicrobat bunyavirus 1 formed distinct clades with vertebrate metagenome-derived sequences, including bats and shrews, indicating either a mammalian origin or the potential to infect mammalian hosts. For example, AusMicrobat phenuivirus 10 (TPM = 0.44) grouped with viral sequences previously identified *Myotis laniger* and *Myotis ricketti* (60% amino acid identity in the RdRp) within the *Phenuiviridae*, strongly suggestive of a bat origin. The remaining viruses (n = 12) were related to invertebrate-associated sequences, including arthropods and protozoan trypanosomatids (**Figure 5A**, **Table 2**).

#### Reoviridae

Twelve reovirus sequences belonging to the family *Sedoreoviridae* were identified, of which nine grouped with viruses previously detected in bat samples, with amino acid identities ranging from 23–92% in the VP1 (RdRp) to their closest relatives (**Figure 5B**). AusMicrobat reoviruses 5–7 fell within the genus *Rotavirus* (TPM = 0.12 – 24,225), grouping with members of the species *Rotavirus A*, *Rotavirus H* and *Rotavirus J*, which are typically mammalian viruses. Of these, AusMicrobat reovirus 6 was found at high abundance in Southern bent-wing bats from the Naracoorte caves, SA (TPM = 24,225), from which we were able to identify 10 out of 11 segments (encoding the VP1, VP3, VP4, VP6, VP7, NSP1, NSP2, NSP3, NSP4 and NSP5 proteins). Similarly, AusMicrobat reoviruses 1-4 clustered with viruses found in metagenome-derived sequences from bats and fish. Conversely, AusMicrobat reoviruses 10–12 were more closely related to invertebrate-associated viral sequences corresponding to unclassified reoviruses.

## Discussion

Bats have long been considered important reservoirs for mammalian related viruses of zoonotic potential [39–41]. Herein, we employed total RNA sequencing on bat faecal samples collected from both natural and artificial roosting sites to identify RNA viral families that are likely to infect mammals [41, 42]. As expected, most of the viruses (∼51.2%) detected were of dietary or parasite (e.g., leishbuviruses) origin, reflecting from the presence of arthropods and plants present in bat droppings [43, 44]. However, despite the small sample size, we also detected a high number of likely mammalian-associated viruses, which seemingly exhibited greater viral richness and abundance than that previously observed in Australian megabats [45–47]. These findings align with those of previous studies that have identified that a broad range of RNA viruses circulate in microbats [41, 44, 48] (**Table 2**, **Figure 1**).

Of particular note was the identification of a novel chuvirus in Australian bats that was closely related to those found in bats and rodents from China, and hence which represents the occurrence of a mammalian-specific chuvirus lineage (as it is untenable that the same dietary-associated viruses would appear in bats on different continents). Although chuviruses were originally described as invertebrate-associated, there is growing evidence for a broader host range [37, 49–51]. Indeed, chuvirus-like sequences have been detected in faecal samples and tissues of bats, rodents, marsupials, and shrews [37, 52], although they did not form a distinct lineage as documented here. The close phylogenetic relationship of AusMicrobat chuvirus 1-2 with Longquan bat chuvirus 1 and Wenzhou rodent chuvirus 1 (∼92 – 95% amino acid identity in the RdRp) strongly suggests that this particular lineage of chuviruses is circulating in mammals on multiple continents (**Figure 3**) [37]. This observation is consistent with the recent isolation of an infectious chu-like virus from tumour cells from the Tasmanian devil (*Sarcophilus harrisii*) indicative of active virus replication in mammalian cells [51] (although this virus lineage is very different to that detected here). More detailed studies of distribution and biology of chuviruses in mammals are therefore clearly warranted.

In a similar fashion to the chuviruses, the clustering of AusMicrobat nodavirus 7 with vertebrate-host derived sequences from Australia, New Zealand and China (**Figure 4**) implies a potential association with bats [53]. Members of the *Nodaviridae* are primarily associated with invertebrates and fish within the *Alphanodavirus* and *Betanodavirus* genera, respectively. However, the discovery of alphanodaviruses with multi-organ distribution such as the porcine nodavirus and bat nodavirus present in brain tissues from pigs and bats, respectively, bolsters the idea that they might also be associated with mammalian infection [37, 54, 55]. Future detection of the novel microbat nodavirus (or its relatives) in bat tissues would provide additional support their association with mammalian infection.

Viruses of likely bat-origin were also identified in a number of other virus families, including the *Coronaviridae, Astroviridae* and *Reoviridae*. With respect to the coronaviruses, we identified bat-associated viruses belonging to three subgenera of *Alphacoronavirus* and related to sequences detected in bats from various geographic locations across Australia, including the Naracoorte caves in South Australia (**Figure 1-2**). The high diversity of astroviruses found in the Naracoorte bat caves suggests that the high population density and close contact of Southern bent-wing bats supports their ongoing transmission and evolution. Indeed, as the Naracoorte bat caves harbour a large maternity colony (∼20,000 to 35,000 individuals), maternal care behaviour might facilitate the circulation of astroviruses from mothers to offspring, particularly given the immature immune systems of newborns and juvenile bats and the extended periods spent together during nursing [56]. These findings align with research on shedding pulses and high levels of astrovirus genetic diversity circulating in cave-roosting bats [57, 58]. Similarly, the detection of viral sequences related to rotaviruses A and J (**Figure 5**) suggests that these cave-dwelling bats harbour a high diversity of rotaviruses.

The data generated here also revealed the geographic connectivity of coronaviruses and caliciviruses between NSW and SA. Of note most, we observed a high diversity and widespread circulation of alphacoronaviruses in microbats within Australia. This was reflected in the phylogenetic clustering of viruses from NSW and South Australia (**Figure 2A-B**), which were similarly closely related to sequences from Queensland, Western Australia and Christmas Island (**Figure 2A-C**). This is indicative of the historical and ecological connectivity of bat populations within Australia and, sporadically, to locations as distant as Christmas Island [19]. In addition, our data provided evidence for virus transmission between bat species. A striking example was the phylogenetic clustering and high sequence similarity of virus sequences found in the little forest bat with those from the large-footed myotis (*Myotis macropus*) and the Christmas Island flying fox (*Pteropus natalis*), suggesting a history of relatively recent cross-species virus transmission [19, 41]. Similarly, the identification of shared caliciviruses between distant roosting locations suggests that microbat populations in the region maintain connectivity through migration, likely sharing seasonal sites, foraging and staging areas (**Figure 1**). Indeed, radio tracking research on southern bent-wing bats from the Naracoorte bat caves showed females flying around 25-35 Km on commuting night flights [59], while monitoring studies of banded bats recaptured Naracoorte bats in Wombeyan, NSW, over 1000 Km apart [60]. Similar findings have been observed for coronaviruses circulating among *Chalinolobus* spp. from Western Australia [19]. Notably, the co-circulation of AusMicrobat calicivirus 1-3 suggest a hidden calicivirus diversity in Lesser long-eared bat populations in SA (**Figure 4**).

In other instances, the data generated here supported far longer virus-host associations, including the possibility of virus-host co-divergence on evolutionary time-scales. This was most apparent in the case of the bat hepeviruses identified here that were most closely related to those previously found in lesser short-tailed bats from New Zealand that diverged from their Australian relatives some 16–18 million years ago [23, 38] (**Figure 3**). Similar findings have been documented for coronaviruses shared between Australian and New Zealand microbats [23]. In addition, the grouping of novel viruses with relatives identified in multiple microbat species was common for the *Astroviridae*, *Rhabdoviridae*, *Coronaviridae* and *Chuviridae*. For instance, virus sequences from the southern bent-wing bat and classified within the *Rhabdoviridae* (AusMicrobat rhabdovirus 3-4) and *Astroviridae* (AusMicrobat astrovirus 10-11), were related to those circulating in *Miniopterus* spp., suggesting that host-jumping and a broader virus host-range might be commonplace [61, 62].

While this study has expanded our knowledge of virus diversity, it says little about their disease potential. As a case in point, although astroviruses are typically associated with gastroenteritis disease in humans, and have been in found microbat skin lesions [47], previous research suggest that these viruses can circulate in apparently healthy bats [63]. As the health status of the sampled Southern bent-wing bat colony in the Naracoorte bat caves is unknown, further studies are needed to determine whether astroviruses represent a pathogenic threat for this critically endangered species [47, 64]. Similarly, relatives of the novel AusMicrobat reoviruses, including Rotavirus A (RVA) and H (RVH), are known to circulate in mammals and birds, including humans, livestock, and bats [65, 66]. Although these cause gastroenteritis disease in some species, it is unclear whether they are pathogenic in bats and the risk they pose for zoonotic transmission [66, 67]. Previous studies have also revealed RVA cross-species transmission events between humans and bats [66, 68, 69]. Of note, *M. schreibersii* has a widespread distribution and a propensity to roost with various other bat species [70], it is important to determine whether the circulation of the novel AusMicrobat rotaviruses poses a potential health threat.

An important limitation of our study was the inability to establish host–virus associations due to the nature of faecal sampling and the limited taxonomic representation of bats in public databases. With the exception of the known presence of the Southern bent wing bat in the Naracoorte caves, the roosting sites were shared by multiple bat species, preventing definitive links between detected viruses and their bat hosts. Although this study offers insights into the likely mammalian-associated RNA virome of microbats, further tissue sampling and follow-up molecular investigations are required to establish host-virus associations, the occurrence of co-infections, and the pathogenic potential of the viruses identified.

This study highlights the high RNA viral diversity in microbats, while the identification of a lineage of mammalian origin in the *Chuviridae* provides an important insight into host-range evolution of these viruses. More broadly, these data further emphasize the ecological and geographical interconnectivity between Australian microbat populations as well as the occurrence of cross-species viral transmission. Future research should address the role of seasonality and roosting habitats in shaping virus circulation and patterns of cross-species transmission among microbats.

## Data availability

The raw sequence reads generated in this study are available at the NCBI SRA database under BioProject PRJNA626677 accessions SAMN49708426 - SAMN49708445 and SAMN49893252 - SAMN49893288. Consensus virus sequences have been deposited in the NCBI/GenBank database under accession numbers XXXX-YYYY.

## Acknowledgements

Data analysis was conducted using the computational resources available on the National Computational Infrastructure (NCI) and the Setonix Supercomputer at the Pawsey Supercomputing Research Centre, Australia. Sampling of microbat faecal samples was made possible through collaboration with Meldanda Reserve–Mid Murray Land care, Naracoorte and Tantanoola Caves, Aldinga Conservation Park, Paxton Wines, and Fairfield council. We are especially grateful to Terry Reardon, Wayne Boardman, Aimee Linke, Don Lester, Ben Paxton, Richard Jasek, Elisa Sparrow, Genki Kondo, Caragh Threlfall, Mei-Ting Kao and Thomas Shortt for their support and collaboration. We also thank Frank Kutsche from the Department for Environment and Water for advice and assistance in facilitating access to sampling locations in South Australia.

## Funding

ECH is supported by a National Health and Medical Research Council (Australia) Investigator grant (GNT2017197).

## Supplementary material

**Supplementary Table S1.** Amino acid substitution models used in the phylogenetic analysis.

## Notes

### Competing Interest Statement

The authors have declared no competing interest.

