## Supplementary Table 1 for "Viral diversity, ecological interconnectedness, and the identification of mammalian chuviruses in Australian microbats"

**Supplementary Table S1.** Amino acid substitution models used in the phylogenetic analysis.

| Group | Model | Protein(s) |
|---|---|---|
| *Astroviridae* | Q.pfam+F+I+$\Gamma_4$ | Capsid |
| *Coronaviridae* | LG+F+I+$\Gamma_4$ \| WAG+F+$\Gamma_4$ | RdRp \| Spike |
| *Rhabdoviridae* | Q.pfam+F+I+ $\Gamma_4$ | RdRp |
| *Bunyavirales* | Q.pfam+F+I+$\Gamma_4$ | RdRp |
| *Caliciviridae* | Q.pfam+F+I+$\Gamma_4$ | RdRp |
| *Chuviridae* | Q.pfam+F+I+$\Gamma_4$ | RdRp |
| *Hepeviridae* | Q.pfam+F+I+$\Gamma_4$ | RdRp |
| *Nodaviridae* | Q.pfam+F+I+$\Gamma_4$ | RdRp |
| *Picornavirales* | Q.pfam+F+I+$\Gamma_4$ | RdRp |
| *Reoviridae* | VT+F+I+$\Gamma_4$ | RdRp |
